# A deep learning approach for uncovering lung cancer immunome patterns

**DOI:** 10.1101/291047

**Authors:** Moritz Hess, Stefan Lenz, Harald Binder

## Abstract

Tumor immune cell infiltration is a well known factor related to survival of cancer patients. This has led to deconvolution approaches that can quantify immune cell proportions for each individual. What is missing, is an approach for modeling joint patterns of different immune cell types. We adapt a deep learning approach, deep Boltzmann machines (DBMs), for modeling immune cell gene expression patterns in lung adenocarcinoma. Specifically, a partially partitioned training approach for dealing with a relatively large number of genes. We also propose a sampling-based approach that smooths the original data according to a trained DBM and can be used for visualization and clustering. The identified clusters can subsequently be judged with respect to association with clinical characteristics, such as tumor stage, providing an external criterion for selecting DBM network architecture and tuning parameters for training. We show that the hidden nodes of the trained networks cannot only be linked to clinical characteristics but also to specific genes, which are the visible nodes of the network. We find that hidden nodes that are linked to tumor stage and survival represent expression of T-cell and mast cell genes among others, probably reflecting specific immune cell infiltration patterns. Thus, DBMs, trained and selected by the proposed approach, might provide a useful tool for extracting immune cell gene expression patterns. In the case of lung adenocarcinomas, these patterns are linked to survival as well as other patient characteristics, which could be useful for uncovering the underlying biology.

## Introduction

The heterogeneity in high dimensional gene expression measurements from tumor specimens partially will be due to different cell types present in the sample. A prominent example is the infiltration of tumors by immune cells which affects patients survival. Immune cell type-specific marker genes have been inferred [1] allowing for techniques such as CIBERSORT [2] that estimate proportions of immune cell types in tumor samples based on gene expression profiles. However these compact representations of immune cell related gene expression lack joint patterns of the abundance of different immune cell types since the representation is limited to discrete estimates of cell type abundances ignoring interactions between cell type-specific marker genes.

Deep learning approaches on the other hand allow for a low-dimensional representation of a large number of measurements by learning the joint distribution of the represented features [3]. Specifically, deep Boltzmann machines (DBMs; [4]) provide a network-structured probability model that can be used for exploring the low-dimensional representation that corresponds to the activation of hidden nodes in the network. The probability distribution learned by the network is manifested in weights between nodes of different abstraction layers that can be actively inspected and interpreted.

Unfortunately, deep learning techniques in general, and DBMs in particular, are limited to settings where the number of individual training data (patients) is much larger compared to the number of features (gene expression). Yet, there are approaches such as partitioning DBMs, which we have recently adapted for genomic data in a different context [5]. Here we adapt such a partitioning approach for modeling immune cell related gene expression in lung adenocarcinoma. Specifically, we propose an approach for selecting between different network architectures and sets of tuning parameters based on visualization and clustering. Subsequently, association with clinical characteristics of patients and specific immune cell groups is investigated. Specifically, the learned representation is then tested for association with tumor stage and patient survival.

In the Methods section, we introduce the lung adenocarcinoma immunome data considered for modeling. Then, we briefly introduce DBMs, including an approach for determining partitions of gene expression features, as a basis for subsequently suggesting different variants of partitioned DBM training to deal with a large number of genes. For judging DBMs, we introduce a sampling-based approach for obtaining a representation of the observed data that is smoothed according to a trained DBM. The latter will be used for clustering the original data, to allow for visually judging DBM model quality based on the resulting patterns. This is illustrated in the Application section. Subsequently, clinical characteristics and different types of immune cell genes will be linked to hidden DBM units. In the Concluding Remarks, we will discuss potential extensions and other promising application areas.

## Materials and methods

### Immune gene type-specific expression data

Gene expression measured in lung adenocarcinoma (LUAD) of 515 different patients was retrieved from the cancer genome atlas (TCGA). Normalized counts were accessed via the Broad Institute TCGA Firehose (data run: July 15th 2016). We considered immune cell type-specific marker genes as provided by Bindea et al. [1]. We removed five genes that showed no variation, resulting in expression measurements for 461 immune cell-type specific marker genes for DBM training.

The range of values observed for different immune cell genes varied widely. This makes it difficult to specify a continuous joint distribution that adequately reflects underlying immune cell proportions. Therefore, we chose to dichotomize the expression for each value at the median across all individuals per gene, in essence using each gene as its own reference. This approach would work best if there were two groups of patients for each cell type, one group with high expression, and the second with low expression, and if the two groups were of equal size. This will probably not be the case, but nevertheless the proposed approach might serve as a good working model in absence of further knowledge.

### Training Deep Boltzmann machines

We model the joint distribution *P* (*X*_1_, …, *X*_*p*_) of *p* dichotomized gene expression measurements by deep Boltzmann machines (DBMs) [4]. Assuming a DBM with two hidden layer (**h**^(1)^, **h**^(2)^), the following log-probability is attributed to the data (**v**):

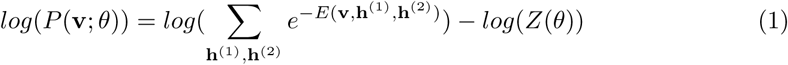

where *θ* corresponds to the parameters of the DBM. *E* is the energy function

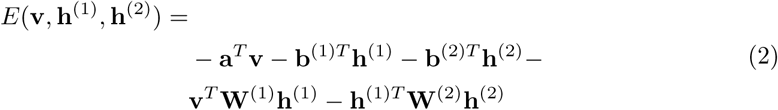

where **W**^(1)^ and **W**^(2)^ are the weight matrices connecting **v** with **h**^(1)^ and **h**^(1)^ with **h**^(2)^ respectively. **a**, **b**^(1)^ and **b**^(2)^ are bias vectors. *log*(*Z*(*θ*)) is the log-partition function

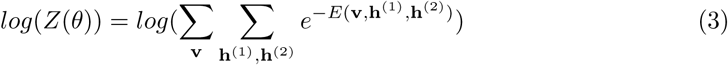

that normalizes the probability. The layered architecture of DBMs enables high-level information to influence parameters in the upper (here second) layer for combining lower-level information for better representation of the data [6].

In order to optimize the likelihood, layer-wise pre-training is employed using contrastive divergence [7], i.e. a two-hidden-layer deep Boltzmann machine is initialized via two stacked restricted Boltzmann machines, RBM1 and RBM2. After the parameters of RBM1 are estimated, RBM2 is trained on the activations of the first layer which are derived by passing the training data through RBM1. Joint refinement of the overall DBM is performed using mean field approximation of the data dependent distribution by variational learning [8] and Gibbs sampling with parallel Gibbs chains for the approximation of the distribution defined by the DBM.

### Partitioned training

To efficiently train a DBM in situations where the number of features (genes) is similar to the number of training data (tumor samples, patients) a partitioned approach was suggested in Hess et al. [5]. Briefly, the idea is to coarsely determine multivariable patterns of dependence by using several multivariable regression model, one for each feature, entering the remaining features as covariates. Based on regression parameters obtained from regularized regression, i.e. on the inferred correlation structure, genes are hierarchically clustered using average linkage. Clusters are obtained by cutting the resulting tree at a specific level such that 10 to 50 genes are clustered together. In Hess et al. [5], a DBM is fully trained for each cluster, and the resulting cluster DBMs are assembled into an overall DBM without further training, setting cross-connections to zero. This allows to consider a number of features that is much larger than the number of observations. In the present setting, the number of features is more moderate. Therefore, we also take cross-connections between cluster DBMs into account.

Specifically, we do not train individual DBMs on each partition but only performed the layer-wise pre-training on different partitions, with refinement performed on the overall, assembled DBM. Furthermore, we allow for flexibility in the training by performing the partitioning in the pre-training only for bottom layers (the first hidden layer in the two-hidden-layer DBM considered subsequently), and adding a joint terminal layer that links the cluster DBMs (Fig 1). This in turn would allow for a better representation of high-level dependencies between gene abundances.

**Figure 1.**
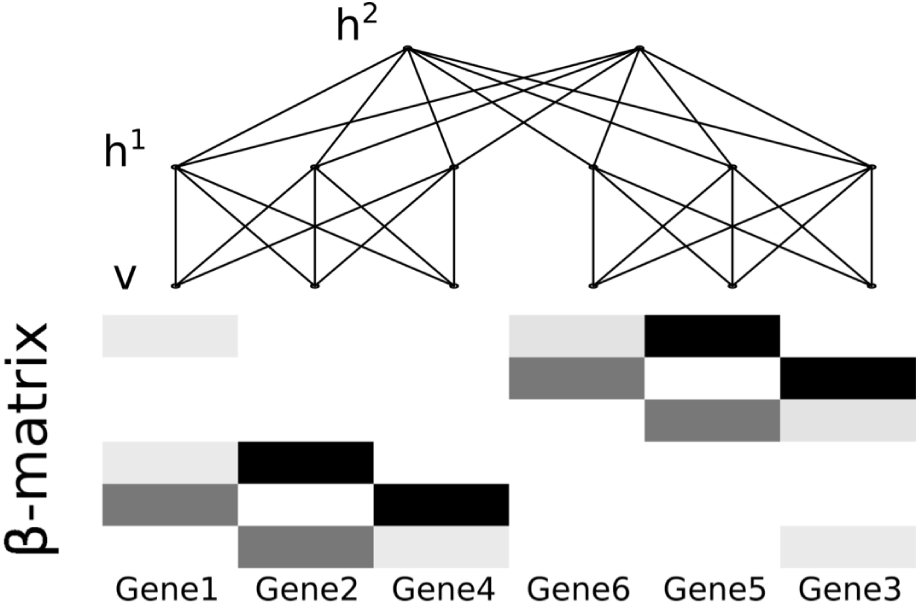
Design of the semi-partitioned DBM. During pre-training, weights between the nodes in the visible layer (v) and the lower hidden layers (only h^1^ in the DBM with two hidden layers, shown here) are initialized separately for cluster of correlated genes. The resulting sub-networks are joined by adding the terminal hidden layer (here h^2^). Correlation between genes is inferred by modeling the mean expression of one gene conditional on the expression of all other genes. Clustering is performed based on the matrix of the resulting *β*s. For details of the clustering see Hess et al. [5]. The design of the semi-partitioned DBM is demonstrated for six hypothetical genes.

Joint refinement of the overall DBM is performed as described above. Since weights that connect the previously partitioned layers in the assembled network are initialized to zero, these weights will receive less adjustment during the joint refinement, effectively leading to regularization of connections between the clusters.

As in Hess et al. [5], we employed a network with two hidden layers. We use *p* nodes in the first layer, and *p/*10 nodes in the second layer to achieve a lower-dimensional representation. For the number of iterations the data is presented to the network (epochs) we considered rather small values, as in Hess et al. [5]. We tested different combinations of the number of epochs for pre-training and joint refinement. For judging the DBMs resulting for different settings, we developed a visualization-based approach, as described in the following.

### Assessing the learned representation based on sampling

We developed a new approach to visualize and evaluate the representation learned by a DBM, based on sampling from the network. The DBM allows to sample from the network by randomly initializing the states of the nodes in the visible and hidden nodes and performing Gibbs sampling for several steps. Yet, in order to arrive at a smoothed representation of the original data, we suggest to initialize the visible units of the network to the original training data. Thereby the Gibbs chain is set to a state close to the original data. After running a Gibbs chain for small number of steps, the visible units correspond to a representation of the original data learned by the network that is still close to the data. We found that 20 steps are sufficient for obtaining such a smoothing effect, and results do not change much with a somewhat larger number of steps, such as 100.

Subsequently, the smoothed representation of the original data is used to infer sample to sample and gene to gene distances, using Euclidean distance. Based on the corresponding distance matrices, patients are clustered using hierarchical clustering and average linkage. While the distances themselves can be used for visualizing DBMs, the clustering result also allows for displaying the original data as a kind of heatmap, sorted according to the smoothed representation. Such a plot can be enriched by clinical annotation, such as tumor stage, to provide a quick glance on whether meaningful patient and gene groups are obtained. To formalize this, we propose to cut the patient clustering tree such as to retrieve two patient clusters, and to inspect for association of these clusters with clinical characteristics. For example, *χ*^2^ statistics will be used when considering association with dichotomized tumor stage, contrasting patients with stage I tumors with stage II, III and stage IV patients.

### Linking hidden units to clinical characteristics and specific immune genes

To extract clinical relevant gene patterns from a DBM, we tested for association between hidden units in the terminal layer (hidden layer two) and the clinical characteristics of interest. Specifically, we propagate activations in the network, starting from the original training data, to obtain for each training sample the activation of nodes in the terminal hidden layer, that can be used in a regression model with the clinical characteristic as outcome. Association between significantly clinically associated hidden nodes and visible nodes (genes), reflecting a strong connection, is investigated analogously. However, since we observed many associations between significant hidden units and visible units, we only considered associations as significant that were stronger than 5% of all the associations between terminal hidden and visible units, observed in the network. The genes corresponding to the resulting visible nodes can be considered to be associated with the clinical characteristics associated with the linked terminal layer hidden unit.

## Results

For adequate modeling of immunome patterns in lung adenocarcinomas, we chose the DBM approach described above. Specifically, we considered expression data from 515 different lung adenocarcinomas to analyze the immune cell related gene expression, represented by 461 immune cell type-specific marker genes.

We used different (partitioned) training approaches and tuning parameters. We set up a standard DBM (DBM), consisting of 461 visible units, 461 units in the first hidden layer and 46 units in the terminal layer. In order to improve learning in presence of a number of gene expression features that is relatively large compared to the number of patients, we also set up different variants of partially partitioned DBMs. In one setting, similar to the approach described in Hess et al. [5], pre-training was performed within completely separated partitions (called “partDBM no joint” in the following). In another setting, as a novel approach, only the visible and first hidden layer were partitioned for pre-training (called “partDBM 1 joint” in the following). In both variants, refinement was performed for the whole DBM. The effect of these choices is illustrated in the following, in particular using the proposed graphical tools.

Clustering the original normalized and dichotomized expression data revealed two patient clusters that demonstrated weak concordance with patient groups differing in tumor stage (Fig 2; original). When using 5 epochs for pre-training and 20 epochs for refinement, clustering the original data based on the learned representations extracted from the network by Gibbs sampling (20 steps) lead to more readily visually apparent structure for the partitioned approaches (Fig 2; partDBM no joint, partDBM 1 joint), while standard DBM training resulted in less structure. Except for the standard DBM, all cluster solutions were significantly associated with tumor stage.

**Figure 2.**
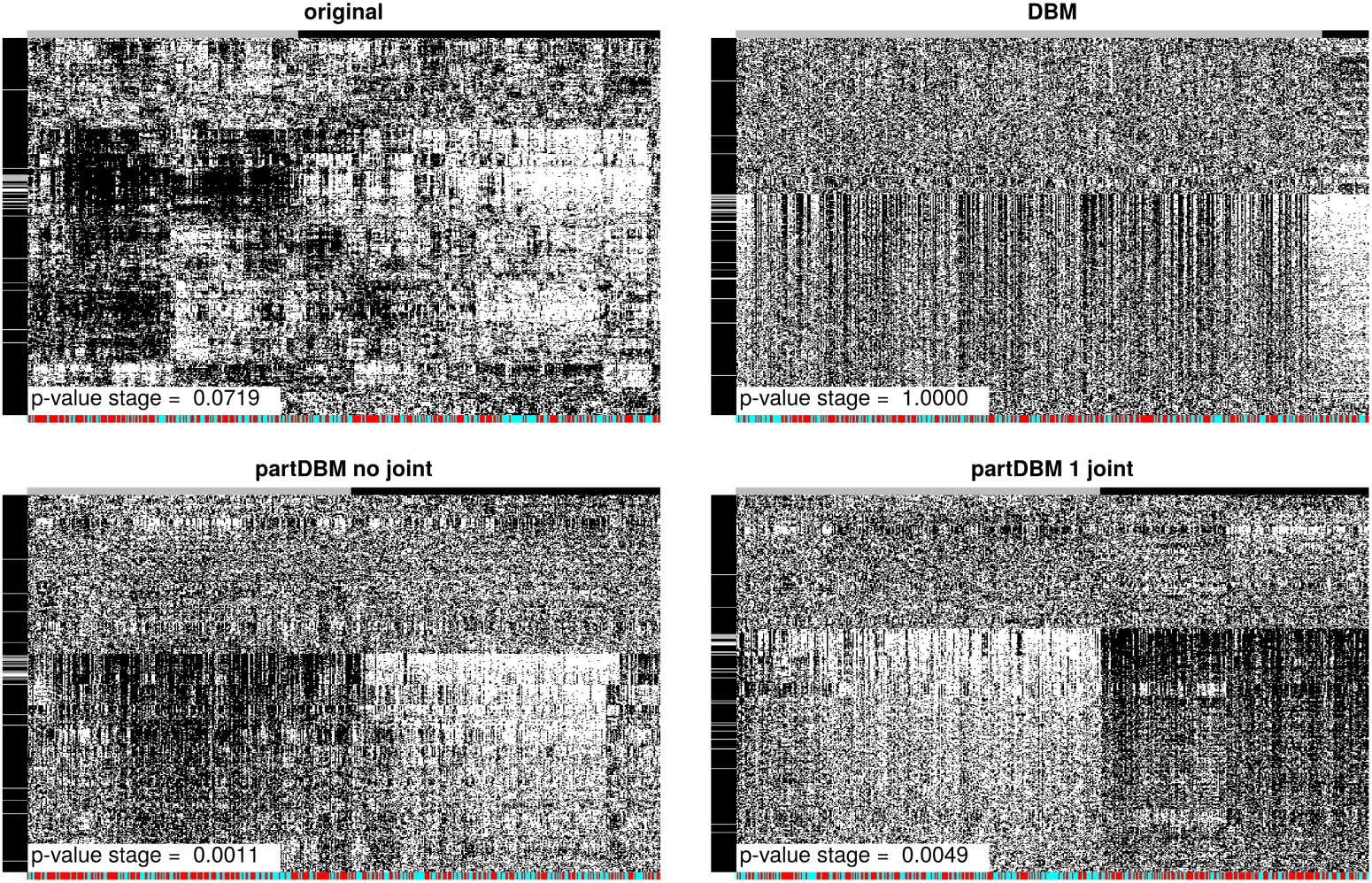
Representations of the lung adenocarcinoma gene expression data learned by DBMs with 5 epochs of pre-training and 20 epochs of refinement. Expression of dichotomized gene expression data of 461 immune cell type-specific marker genes in 515 tumors is shown. Expression was either clustered based on distances inferred from the original data (original) or based on distances inferred from the learned representations of the DBM. The learned representations were extracted by sampling from the network after running a Gibbs chain for 20 steps (DBM, partDBM no joint, partDBM 1 joint). White and grey horizontal lines in the left margin bar indicate T-cell and Cytotoxic cell type-specific marker genes, respectively. Solutions with two patient clusters inferred from clustering the original expression data or the learned representations are indicated by vertical gray and black colored bars respectively. Tumor stage groups are indicated by cyan and red colored vertical bars in the bottom margins. Concordance of patient groups and inferred patient clusters is determined by a *χ*^2^ test.

Yet, the association with tumor stage strongly depends on the number of epochs used for training, besides the overall training approach, as seen from Fig 3. In particular the unpartitioned DBM did only perform well in a certain combination with a very large number of epochs, which might be prone to overfitting. The DBMs which were partially partitioned performed better in many scenarios and were more robust against variation of the number of epochs. The partially partitioned DBM where only the visible and the first hidden layer were partitioned during pre-training (“partDBM 1 joint”) seems to perform best with an intermediate number of epochs, with performance degradation for a larger number of epochs. In contrast the DBM that was fully partitioned during pre-training (“partDBM no joint”) might even result in better results for a larger number of epochs. Overall, using 5 epochs for pre-training and 20 epochs for joint-refinement, i.e. the setting shown in Fig 2, led to good concordance between clusters extracted from the learned representation and patient groups differing in tumor stage.

**Figure 3.**
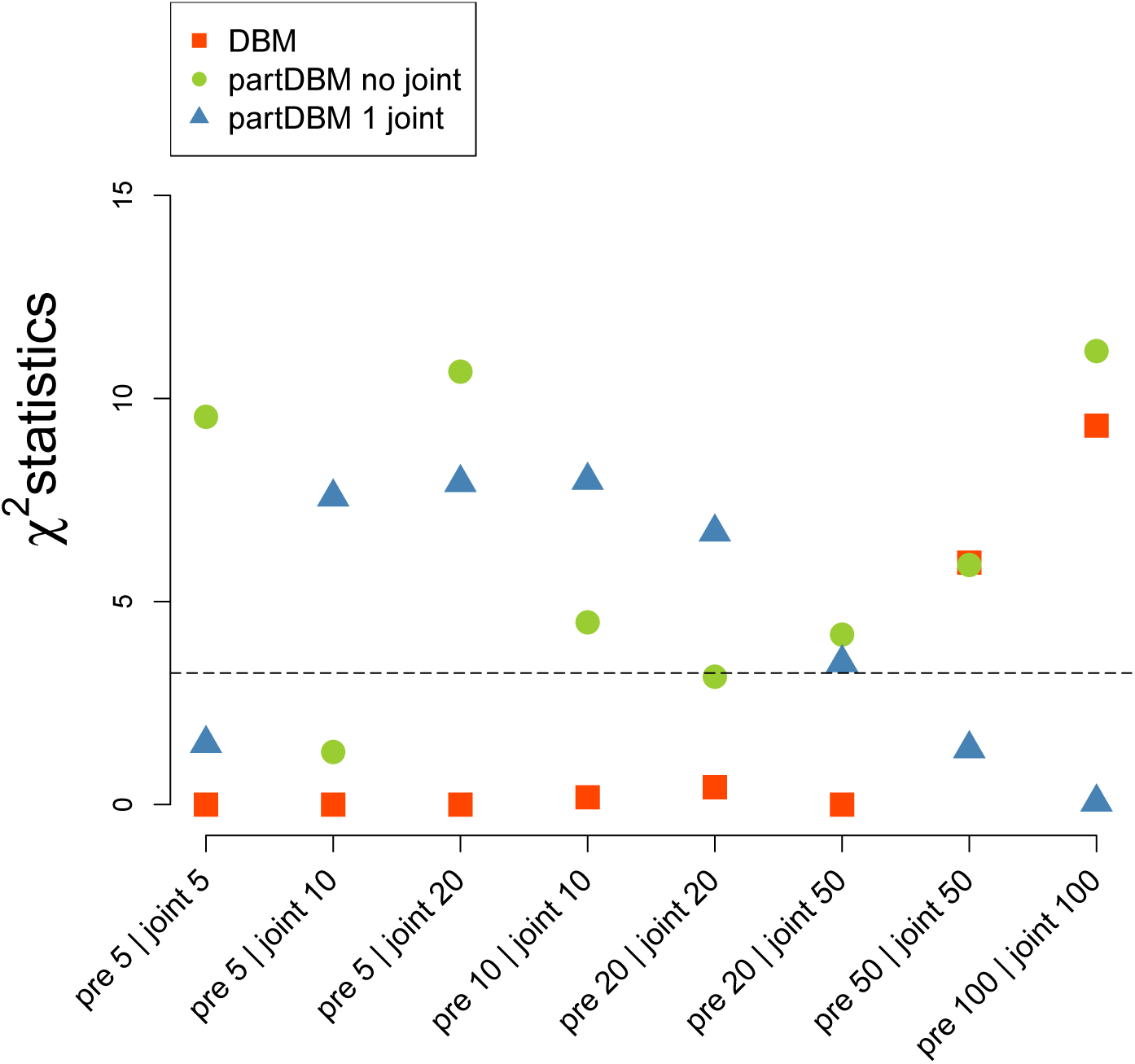
Concordance of patient groups that differ in tumor stage with patient clusters, inferred from the expression data using DBMs. Three different network architectures (DBM, partDBM no joint and partDBM 1 joint) were evaluated using eight different combinations of epochs during which layer-wise pretraining (pre) or joint refinement (joint) was performed. Concordance among identified patient cluster and previously defined patient groups (stage I patients vs. stage II, III and IV patients) was assessed by *χ*^2^ statistics. The horizontal line represents the concordance observed when clustering the original representation of the gene expression data.

Although the three architectures differed in the performance to learn the differential gene expression patterns, observed between patients differing in tumor stage, the DBM training approaches performed equally well in clustering genes into a group that does not contribute to the patient clustering and a group of genes that seemed to drive patient clustering. This is indicated by a group of immune cell type-specific gene categories (T cells and Cytotoxic cells) that were similarly clustered in all three network architectures (white and gray horizontal lines in the left border bars of Fig 2).

Having assured association of the patterns that were learned by the DBMs with tumor stage, we also investigated association with the clinical endpoint survival as well as with specific gene groups, to obtain a more complete picture of the underlying biological process. Correspondingly, we tested the terminal hidden units for association with tumor stage as well as for association with survival, using Cox proportional hazards models for the latter. Interestingly we found several nodes that were both connected to survival and tumor stage. We exemplarily selected two of the most strongly associated hidden nodes and inferred their connections to immune cell marker gene categories. One hidden node (Fig 4; 1) was strongly connected to T-cell, Cytotoxic cells and B-cell marker gene expression while the other node was rather connected with Mast cells or TFH cells. Thus different cell type groups might be associated jointly with tumor stage and patient survival.

**Figure 4.**
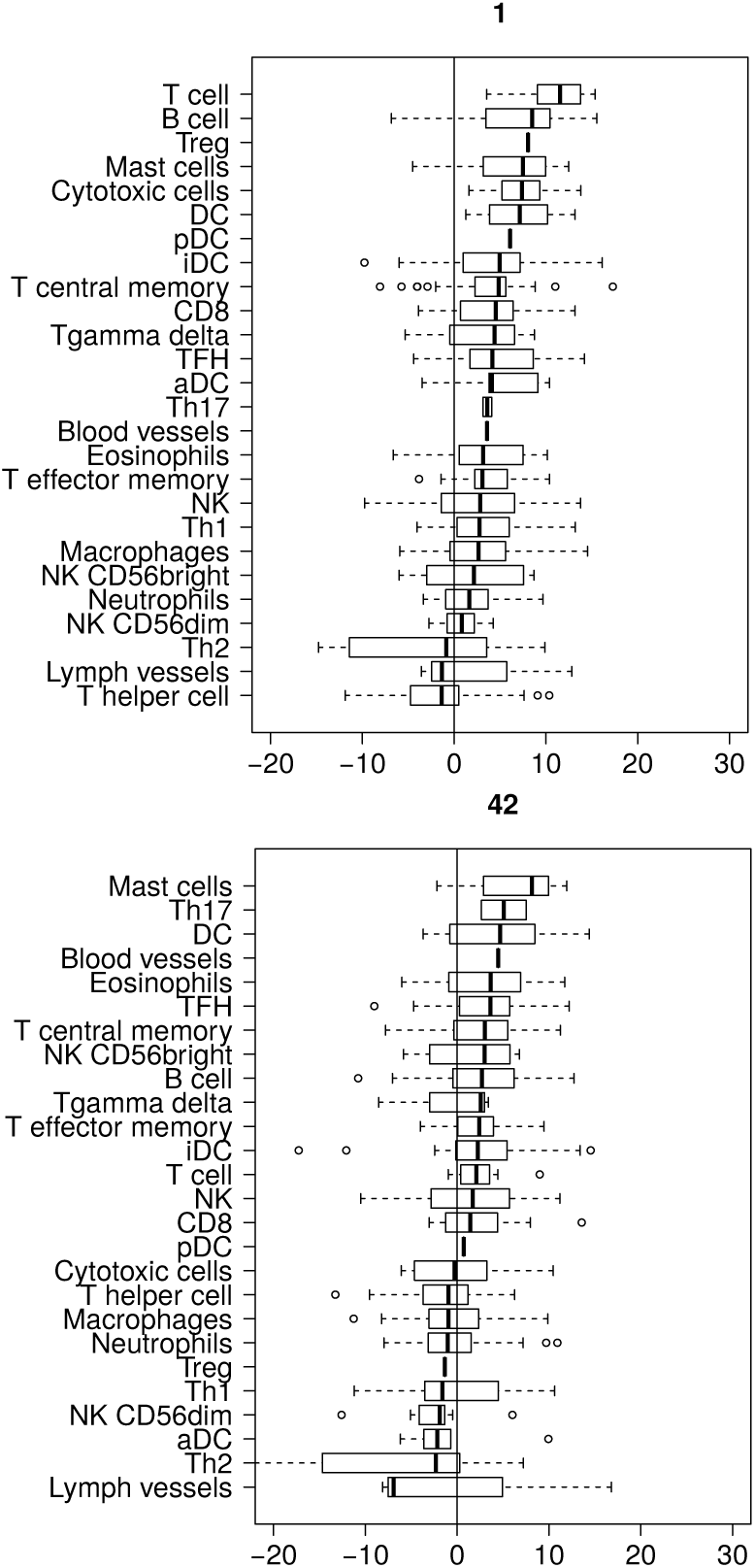
Connection of hidden nodes, that were associated with tumor stage and survival, with immune cell type-specific marker gene categories. Significant hidden nodes (FWER ≤ 0.05) were tested for association with visible units (genes) using linear regression models. Connections between terminal hidden units and visible units (genes) were considered significant if they were stronger than 5% of all connections found between the visible layer and the terminal hidden layer. The boxes indicate the distribution of z-scores that indicate the strength of connection of the respective marker-gene group with the hidden nodes. The IDs of the hidden nodes are indicated in the figure headings.

## Discussion

The immunome in cancer can be investigated by considering the expression levels of immune cell type-specific genes. We investigated complex patterns in such expression levels for lung adenocarcinoma using deep Boltzmann machines (DBMs). To be able to do so we adapted a partitioned learning approach that can deal with a number of genes that is relatively larger compared to the number of patients, in contrast to many other deep learning approaches.

Specifically, we investigated a flexible extension of the partitioning approach presented in Hess et al. [5]. We compared the performance of the approach with unpartitioned deep Boltzmann Machines and evaluated tuning parameter settings. To be able to select a good deep Boltzmann machine, we introduced a visualization-based approach. This relied on a smoothed version of the original data, moved closer to the DBM network representation.

Heatmap-type plots of the original data, sorted according to the smoothed version, allowed to visually assess association of identified gene and patient groups with clinical characteristics. This was formalized via a two-cluster-based criterion and served to select a DBM for subsequent more detailed analysis. Specifically, the DBMs obtained from partitioned training were superior compared to unpartitioned training. These results suggest that regularization, here performed by constraining the weights connecting weakly correlated genes, does allow for improved learning of a meaningful representation when the number of investigated features is equally large compared to the number of independent training data. As a consequence, deep Boltzmann machines may be applied in many scenarios occurring in the field of biostatistics in order to explore the data and to extract interpretable patterns. In our example, terminal layer hidden nodes of DBMs trained by the newly proposed approach were found to be associated with tumor stage and survival. These nodes could be linked to genes that reflected specific immune cells, such as T-cells, Cytotoxic cells or mast cells. Tumor infiltration by cytotoxic cells is associated with improved survival in lung cancer patients [9] and a high amount of T-cells is generally known to be linked with good prognosis in many cancers including lung cancer [10]. This concordance suggests biological relevance of our findings which sheds some light on potential mechanisms and demonstrates the power of deep Boltzmann machines to unravel patterns in the data. Since we observed a partially differential performance between our partitioned training approaches, we plan to further investigate which partitioning scheme performs well in which particular scenario, in order to optimize the training.

## Conclusion

Deep Boltzmann machines are a promising approach for learning compact representations of high dimensional data such as gene expression data. Using immune cell type-specific gene expression in lung adenocarcinoma as model, we evaluated a new approach for partially training partitioned DBMs on subsets of correlated features and selecting between DBMs obtained from different training schemes. The resulting DBM could be linked to clinical covariates as well as specific immune cell types, which shows that properly trained DBMs can be useful for gaining biological insight, beyond black box prediction applications that are dominant in the deep learning field so far.

## Acknowledgments

This work has been supported by the project “Intrinsische Strahlenempfindlichkeit: Identifikation biologischer und epidemiologischer Langzeitfolgen” (ISIBELA) funded by the BMBF, Fkz. 02NUK042 and by the project “Transfer of cognitive training gains in cognitively healthy aging: Mechanisms and Modulators” (AgeGain) Fkz. 01GQ1425A. The results shown here are in part based upon data generated by the TCGA Research Network: http://cancergenome.nih.gov/.

